# Morphology of petiole bending, senescence, epinasty, along with necrotic scarring in tomato leaves infiltrated with virulent *Ralstonia pseudosolanacearum*

**DOI:** 10.64898/2026.05.21.711296

**Authors:** Monika Jain, Sumi Kalita, Priya Rani Daimari, Zoomani Rabha, Shuhada Begum, Lukapriya Dutta, Shubhra Jyoti Giri, Shuvam Bhuyan, Sunita Kushwah, Aditya Kumar, Suvendra Kumar Ray

**Affiliations:** Department of Molecular Biology & Biotechnology, Tezpur University, Tezpur–784028, Assam, India

**Keywords:** Leaf infiltration, *Ralstonia pseudosolanacearum*, Petiole bending, Necrotic scar, Senescence, Epinasty

## Abstract

*Ralstonia pseudosolanacearum* (*Rps*) belongs to the *Ralstonia solanacearum* species complex (RSSC). It is a vascular pathogen that causes lethal bacterial wilt disease in many plants, including tomato and eggplant. In this study, we infiltrated tomato leaves with the phytopathogenic bacterium at 10^9^ CFU/mL and observed the development of necrotic scars in the infiltrated area at 48 hours post-infiltration. Interestingly, this response was followed by petiole bending toward the ground of the compound leaf. This was followed by the gradual senescence of the infiltrated leaflet only. In addition, the terminal leaflet infiltrated with the pathogen exhibited epinasty. None of the above symptoms were observed in leaves infiltrated with the known virulent deficient *hrpB::*Ω mutant. Surprisingly, all of the above symptoms were observed in leaves infiltrated with another well-known virulence-deficient mutant *phcA::*Ω. It indicated that the necrotic lesion caused in tomato leaves was *hrp*-dependent. Infiltration in eggplant leaves caused necrotic scarring and leaf senescence, which were relatively delayed. Necrotic scarring without petiole bending or senescence in tomato leaves was also observed due to infiltration of *Pseudomonas aeruginosa* SPT08, a tomato endophyte having plant growth promotion activity. The patho-phenotypes such as petiole bending, epinasty, and senescence observed in the case of tomato in this study were not reported earlier. We believe these phenotypes produced in tomato after leaf infiltration may be useful to study the virulence of this pathogen.

## 1 Introduction

*Ralstonia solanacearum* Species Complex (RSSC) is among the most devastating plant pathogenic bacteria worldwide (Vailleau and Genin 2023). It causes a lethal wilt disease infecting a range of hosts, including common fruits and vegetables such as eggplant, chilli, potato, and tomato (Lowe-Power et al. 2020). The pathogen is widely distributed across diverse geographical regions, including America, Africa, and Asia. It is one of the oldest plant pathogenic bacteria to be discovered by researchers (Hayward 1991), which might be due to its preponderance and economic importance in the world. The pathogen naturally enters the host plant through the root and systemically colonizes the xylem vessels of the entire plant, ultimately occluding the water-transport system and causing wilting and death of the host (Hayashi et al. 2019; Schell 2000; Genin 2010).

Researchers worldwide adopt the classic *in vivo* soil drenching approach as a pathogenicity assay in the grown-up tomato plants, as it closely resonates with the natural mode of soil-borne infection (Giri et al. 2025). It takes at least two weeks after inoculation to complete the virulence assessment of a strain using the soil inoculation method. Some challenges in this approach are inconsistency in observing the wilting symptoms and the frequent occurrences of escapees (Bhuyan et al. 2025b), which are plants systemically colonized by the pathogen but are completely asymptomatic. This unpredictable and inconsistent behavior often extends the virulence assessment more than four weeks post-inoculation. Methods such as stem inoculation (Diatloff and Imrie 2000; Johnson and Cummings 2013; Morel et al. 2017) and petiole inoculation (Fry and Milholland 1990; Saile et al. 1997; Thomas et al. 2015; Khokhani et al. 2018) in grown-up plants have been developed to study its virulence functions. Recently, inoculations in both the root and the cotyledon leaves of the seedlings have been developed to study virulence functions (Kumar et al. 2013; Kumar et al. 2017; Singh et al. 2018; Bhuyan et al. 2026).

It is common but surprising that a virulent *Ralstonia pseudosolanacearum* (*Rps*) strain can colonize a susceptible tomato seedling or plant without observation of disease symptoms in the plant, which is a frequent observation by the researchers studying bacterial wilt (Bhuyan et al. 2025b, Bhuyan et al. 2026). It might be a profound evolutionary adaptation for this systemic pathogen. Considering disease development is a complex process governed by multiple factors as described by the plant disease triangle (Fletcher et al. 2013), development of an assay that is both faster and capable of unambiguously demonstrating virulence is very much needed in bacterial wilt research.

In this study, we have observed necrotic lesions in tomato leaves 48 h post infiltration with *Rps*. Apart from the necrotic lesions, several other phenotypes such as petiole bending, senescence, and epinasty have been observed in tomato leaves infiltrated with pathogen. These pathophenotypes may be used in future to study virulence functions of *Rps*.

## 2 Materials and Methods

### 2.1. Bacterial strains and growth conditions

Bacterial strains used in this study are listed in Table 1. *Ralstonia pseudosolanacearum* F1C1, its derivative mutants (*hrpB::*Ω and *phcA::*Ω), and *Pseudomonas aeruginosa* SPT08 were cultured in BG broth composed of 0.1 % yeast extract (Hi-Media, Mumbai, India), 1 % peptone (Hi-Media), 0.1 % casamino acids (SRL, New Mumbai, India), and supplemented with 0.5 % glucose (Hi-Media) at 28 °C (Bhuyan et al. 2023). *Escherichia coli* DH5α was grown in 2% LB medium (Hi-Media) at 37 °C. *Xanthomonas oryzae* pv. *oryzae* (*Xoo*) BXO43 and its derivative mutant defective in Type III secretion system (T3S) were grown in PS (Peptone-Sucrose) media containing 1% peptone and 1% sucrose (Hi-Media) at 28 °C (Ray, 2001). Solid media were prepared by supplementing the respective broths with 1.5% agar (Hi-Media). The Rifampicin (Hi-Media), Spectinomycin (Hi-Media) and Kanamycin (Hi-Media) antibiotics were used at 50 μg/mL.

**Table 1.** Bacterial strains used in the study.

| Strains | Characteristics | Source |
| --- | --- | --- |
| *Ralstonia pseudosolanacearum* |  |  |
| F1C1 | Wildtype Vir<sup>+</sup>, EPS<sup>+</sup> strain belonging to phylotype I | Kumar et al. (2013) |
| *phcA::Ω* | Vir<sup>-</sup>, EPS<sup>-</sup>, Spc<sup>r</sup> mutant derived from F1C1 after Ω interposon insertion | Phukan et al. (2019) |
|  | in *phcA* locus in the genome |  |
| *hrpB*::$\Omega$ | Vir<sup>-</sup>, EPS<sup>+</sup>, Spc<sup>r</sup> mutant derived from F1C1 after $\Omega$ interposon insertion in *hrpB* locus in the genome | Phukan et al. (2019) |
| *Escherichia coli* |  |  |
| DH5α | F-Φ80*lacZ*ΔM15 $\Delta$ (*lacZYA-argF*) U169 *recA1 endA1 hsdR17* (rK<sup>-</sup>, mK<sup>+</sup>) *phoA supE44 λ-thi-1 gyrA96 relA1* | Laboratory collection |
| *Pseudomonas aeruginosa* |  |  |
| SPT08 | Wildtype plant growth-promoting bacteria isolated from tomato seedlings that exhibits antagonistic activity against F1C1 | Giri et al. (2026) |

### 2.2. Seed germination and growth of mature plants

Seeds of tomato (cv. Pusa Ruby; Durga Seeds, U.P., India) and eggplant (cv. Dev-Kiran (614); Tokita, Karnataka, India) were used in this study. Both tomato and eggplant seeds were germinated according to the protocols outlined by Bhuyan et al. (2025b) and (2026). Prior to sowing, tomato seeds were soaked in sterile dH_2_O at room temperature for 24 h, whereas eggplant seeds were soaked in sterile dH_2_O at 4 °C for 48 h. The seeds were then sown in containers containing sterile soil, sealed in zip-lock plastic bags to retain moisture, and kept in the dark at ambient temperature for 72 h. After germination, seedlings were transferred to a growth chamber (Orbitek, Scigenics, India) maintained at 28 °C with a 12 h light/dark cycle and approximately 80% relative humidity. The seedlings were watered at regular intervals to prevent desiccation. Under these conditions, the majority of seeds germinated within 7 days, reaching the cotyledon stage.

The 7-day-old cotyledon stage seedlings were then transplanted into 500 mL reusable plastic pots containing sterile soil and maintained under greenhouse conditions at 28 °C with a 12 h light/dark cycle. Plants were watered at regular intervals to ensure adequate growth and were maintained for two months prior to experimental use.

### 2.3. Preparation of bacterial inoculums

Freshly cultured bacterial cells were introduced into 10 mL of BG broth and incubated in a shaking incubator (Orbitek, Scigenics, India) under optimal temperature conditions for each strain at 150 rpm for overnight. After incubation, cells were harvested by centrifugation at 4000 rpm for 15 min at 4 °C, and the resulting pellet was resuspended in an equal volume of sterile dH_2_O. The cell density was adjusted to approximately 10 CFU/mL (OD ≈ 1.0). For subsequent experiments, 10-fold dilutions of the 10^9^ CFU/mL were prepared using the serial dilution method till 10^3^ CFU/mL.

### 2.4. Infiltration of leaves with bacterial inoculums

The washed cell suspension was drawn into a sterile syringe. After removing the needle, the nozzle was gently pressed against the dorsal (abaxial) surface of a fully grown plant leaf. Infiltration was achieved by applying mild pressure to allow the suspension to enter the leaf tissue through the stomata, following the method as illustrated in Fig. 1. The infiltration was performed in different leaflets of a compound leaf in tomato or different leaves separately in the plant. The procedure was followed for strains such as wild-type *Rps* F1C1 (Kumar et al 2013), F1C1-derived virulence-deficient mutants such as *hrpB::*Ω and *phcA::*Ω (Phukan et al 2019), a known endophyte *Pseudomonas aeruginosa* SPT08 isolated in our laboratory (Giri et al 2025), and *Escherichia coli* DH5□, a non-phytopathogenic control. Sterile dH_2_O was also infiltrated as negative control. The infiltration was also repeated for *Xanthomonas oryzae* pv. *oryzae* strain BXO43 which is reported to give a hypersensitive response (HR) in non-host tomato, and the BXO43 derived mutant defective in T3S which is HR^−^ in tomato (Jha et al., 2007) to understand how the two responses differ. In the case of eggplant, which bears simple leaves, infiltration was performed on different leaves of a plant. The infiltrated leaves were monitored afterwards for any visible responses.

**Figure 1.**
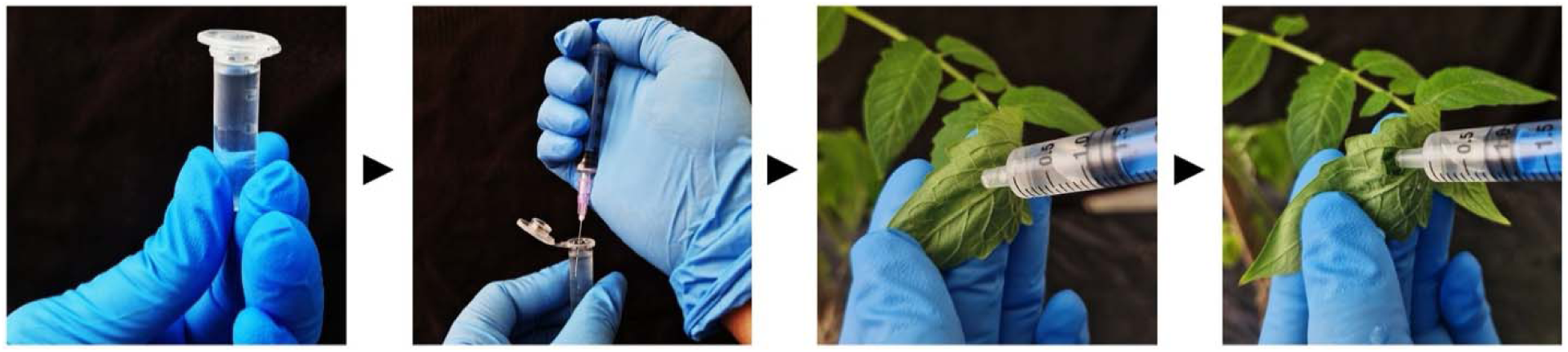
Infiltration of bacterial suspension into the dorsal side of the tomato leaf.

## 3 Results

### 3.1 Observation of necrotic scars in the infiltrated leaflet of tomato plant that gradually led to senescence

Leaf infiltration of the pathogen resulted in the development of a necrotic scar in the infiltrated area within 48 hours post-infiltration in the case of tomato (Fig. 2). At different pathogen concentrations, ranging from 10^9^ to 10^6^ CFU/mL, a distinct necrotic scar formed, limited to the infiltrated region of the leaf by 72 h after infiltration. The scar remained limited to the infiltrated area and did not spread to the adjacent leaflet within a compound leaf even after 10 days post-infiltration (DPI), however, the affected leaflet gradually exhibited yellowing, indicative of senescence (Fig. 3). To determine whether scar formation in the leaf was associated with *Rps* virulence function, we used an F1C1-derived *hrpB::*Ω mutant lacking a functional type III secretion system (T3S), which is avirulent in both tomato seedlings and mature plants, and fails to elicit HR in tobacco leaves (Phukan et al. 2019). The infiltration of F1C1-derived *hrpB::*Ω mutant failed to develop any scar, and the infiltrated leaf appeared similar to those mock-infiltrated with sterile water or with the non-phytopathogenic *E. coli* DH5α (Fig. 4). Likewise, infiltration with *Escherichia coli* into tomato leaves did not result in any scar formation up to 10 DPI. Further, the F1C1-derived *phcA::*Ω mutant, which was a known virulence-deficient mutant (Phukan et al., 2019), when infiltrated into tomato leaves, developed a scar like those caused by the wild type. It suggested that leaf-infiltration might serve as a sensitive method to study virulence functions of this pathogen. It also suggests that mutants that exhibit virulence deficiency by root inoculation may not be virulence-deficient by direct cell-to-cell interaction with plant cells.

**Figure 2.**
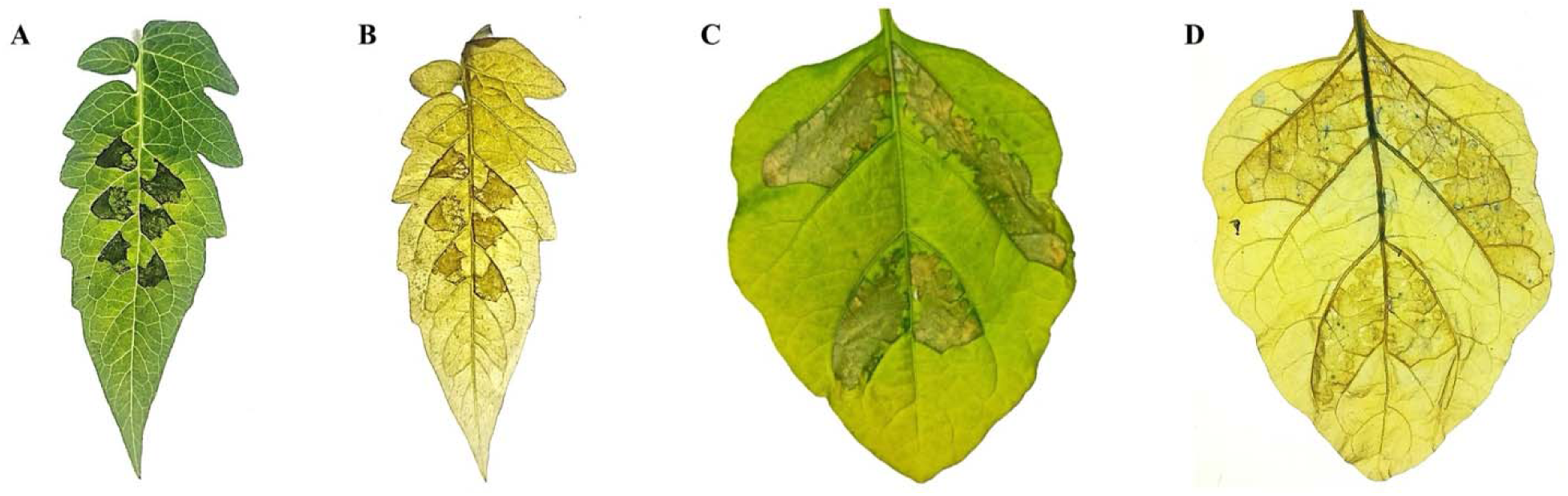
Localized necrotic scarring on infiltrated leaves. **[A]** Tomato leaf at 48 h, and **[C]** eggplant leaf at 96 h following bacterial infiltration. **[B & D]** Corresponding trypan blue-stained tomato and eggplant leaves highlighting plant tissue death in the infiltrated region.

**Figure 3.**
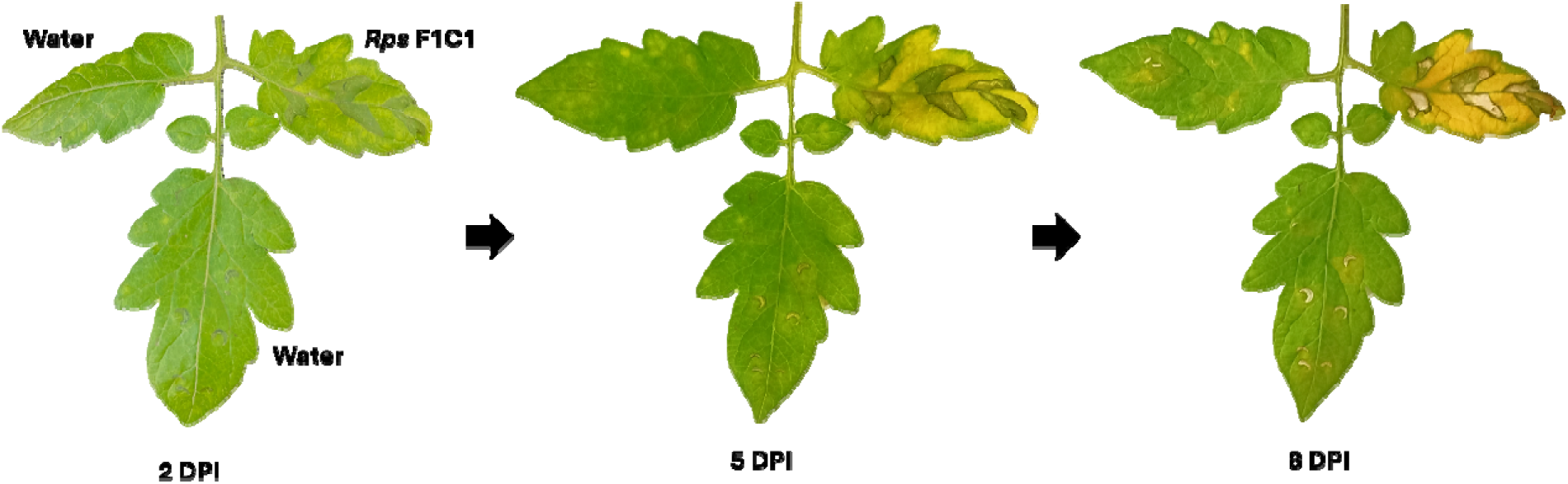
The symptom of necrotic scar and senescence remains confined to the pathogen-infiltrated leaflet and the adjacent water infiltrated-leaflet remains unaffected.

**Figure 4.**
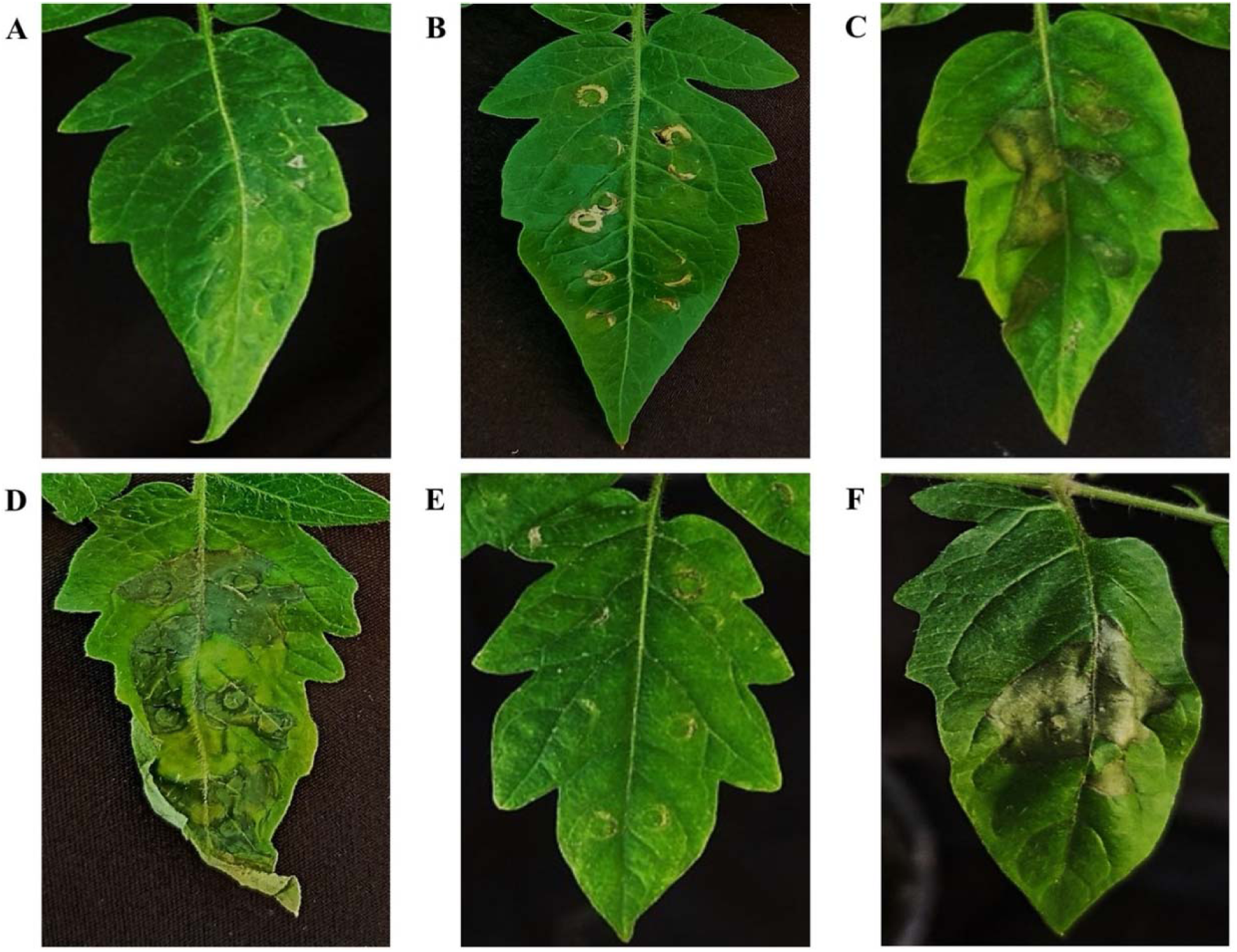
Tomato leaflets exhibiting necrotic scarring at 5 DPI. **[A]** dH_2_O control; **[B]** *E. coli* DH5□; **[C]** *Rps* F1C1; **[D]** *phcA::*Ω; **[E]** *hrpB::*Ω; **[F]** *P. aeruginosa* SPT08. This experiment was independently performed by four researchers with multiple biological replicates.

### 3.2 Observation of petiole bending and epinasty of the terminal leaflet of tomato post leaf-infiltration

After 48 hours of leaf infiltration, besides the developed scar, the compound leaves of tomato exhibited downward bending at the petiole. Under normal conditions, the compound leaf petiole forms an acute angle (∼65 °) with the plant’s vertical axis (Fig. 5); however, following infiltration, bending occurred, and this angle progressively increased to become obtuse, starting at 3 DPI. This response of petiole bending was observed in leaves infiltrated with wild-type F1C1 and *phcA::*Ω, but not with *hrpB::*Ω or water or *E. coli* DH5α infiltration, which also failed to induce the symptom of necrotic scarring. Notably, petiole bending was detectable even at a low bacterial concentration of 10^5^ CFU/mL, at which no visible scar was observed, underscoring the sensitivity of the assay.

**Figure 5.**
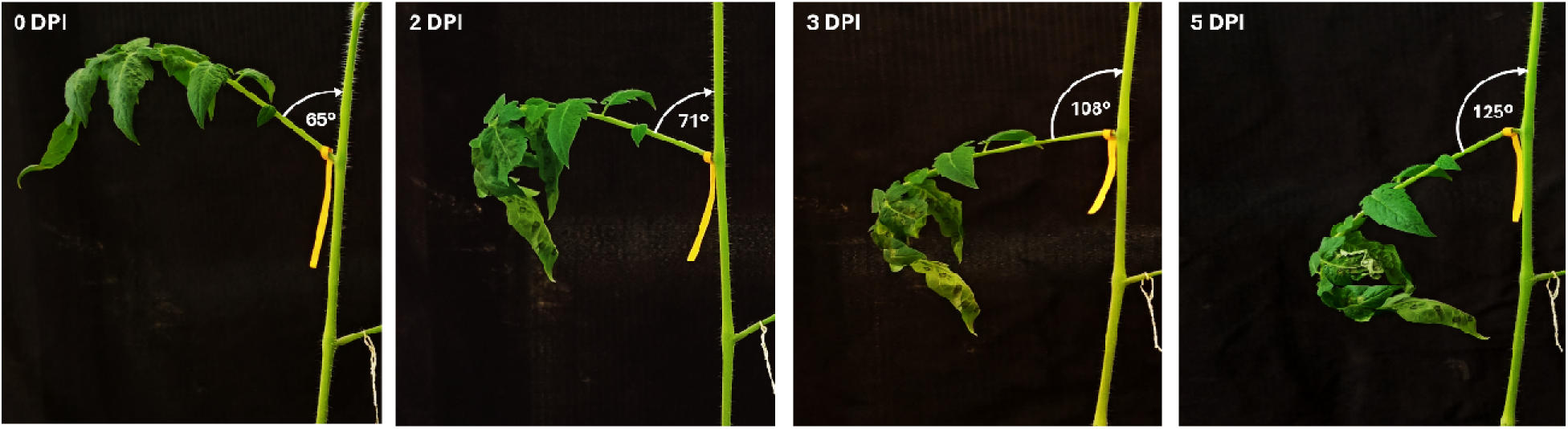
Progression of petiole bending in tomato leaves following bacterial infiltration. Epinasty of the terminal leaflet was evident since 3 DPI (72 h post-infiltration). This experiment was independently performed by four researchers with multiple biological replicates.

Furthermore, 72 hours post-infiltration, leaves infiltrated with either wild-type F1C1 or *phcA::*Ω, displayed pronounced epinasty in the terminal leaflet of the tomato (Fig. 4). According to existing literature, these pathogenicity features have not been reported previously.

### 3.3 Observation of necrotic scars in eggplant leaf post *Rps* infiltration

Mature eggplants were also subjected to leaf infiltration with the cells of F1C1, *hrpB::*Ω, *phcA::*Ω, and SPT08 (Fig. 6). In eggplant leaves, a necrotic scar formation wa observed following infiltration with F1C1 (Fig. 7) and *phcA::*Ω, although with a delayed onset (∼3 DPI) compared to tomato. In contrast, the *hrpB::*Ω mutant did not induce any symptoms, consistent with its behavior in tomato. Interestingly, SPT08 did not induce necrotic scarring in eggplant leaf (Fig. 6), unlike its response in tomato (Fig. 4).

**Figure 6.**
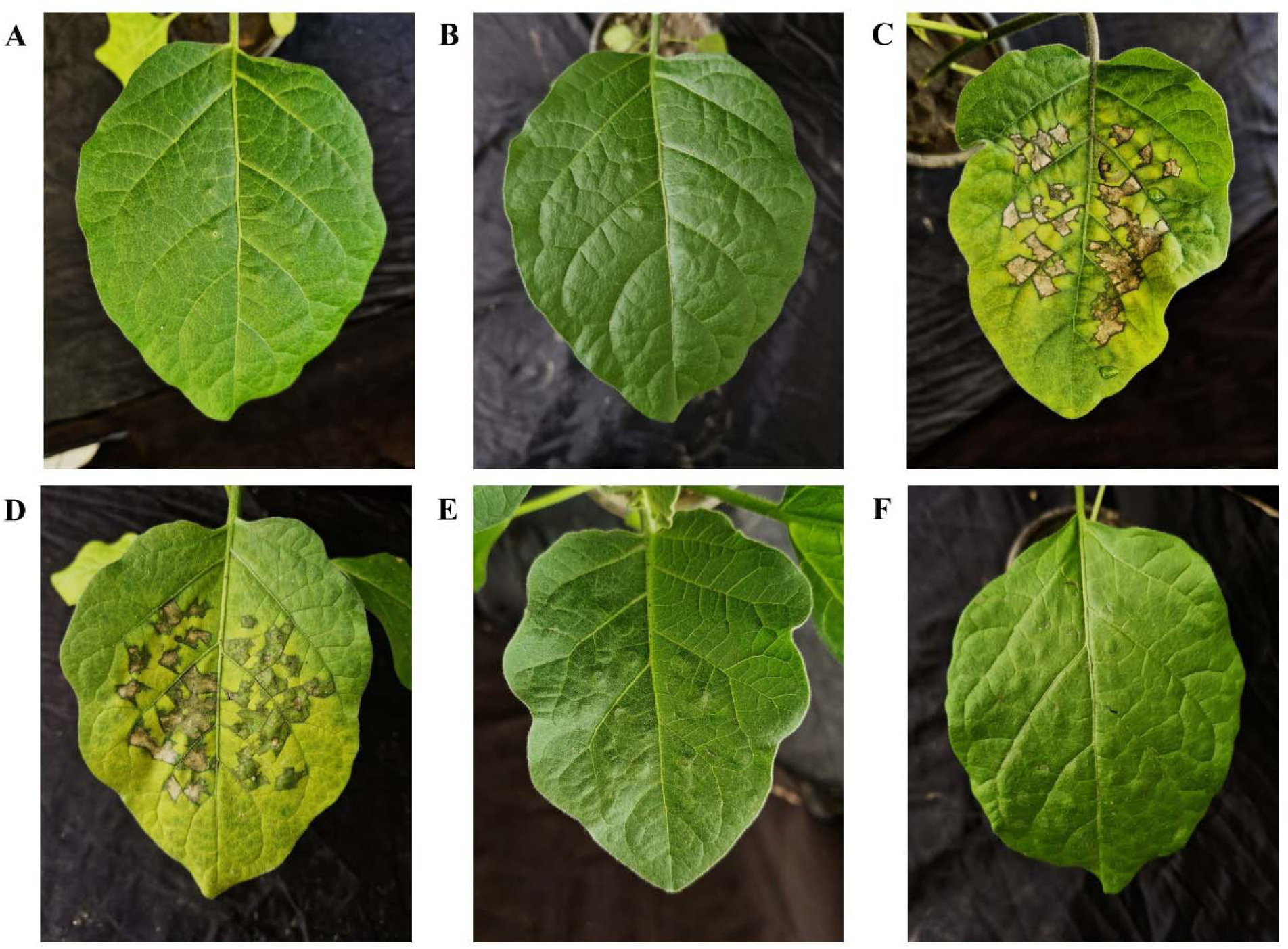
Eggplant leaves exhibiting necrotic scarring at 6 DPI. **[A]** dH_2_O control; **[B]** *E. coli* DH5□; **[C]** *Rps* F1C1; **[D]** *phcA::*Ω; **[E]** *hrpB::*Ω; **[F]** *P. aeruginosa* SPT08. This experiment was independently performed by four researchers with multiple biological replicates.

**Figure 7.**
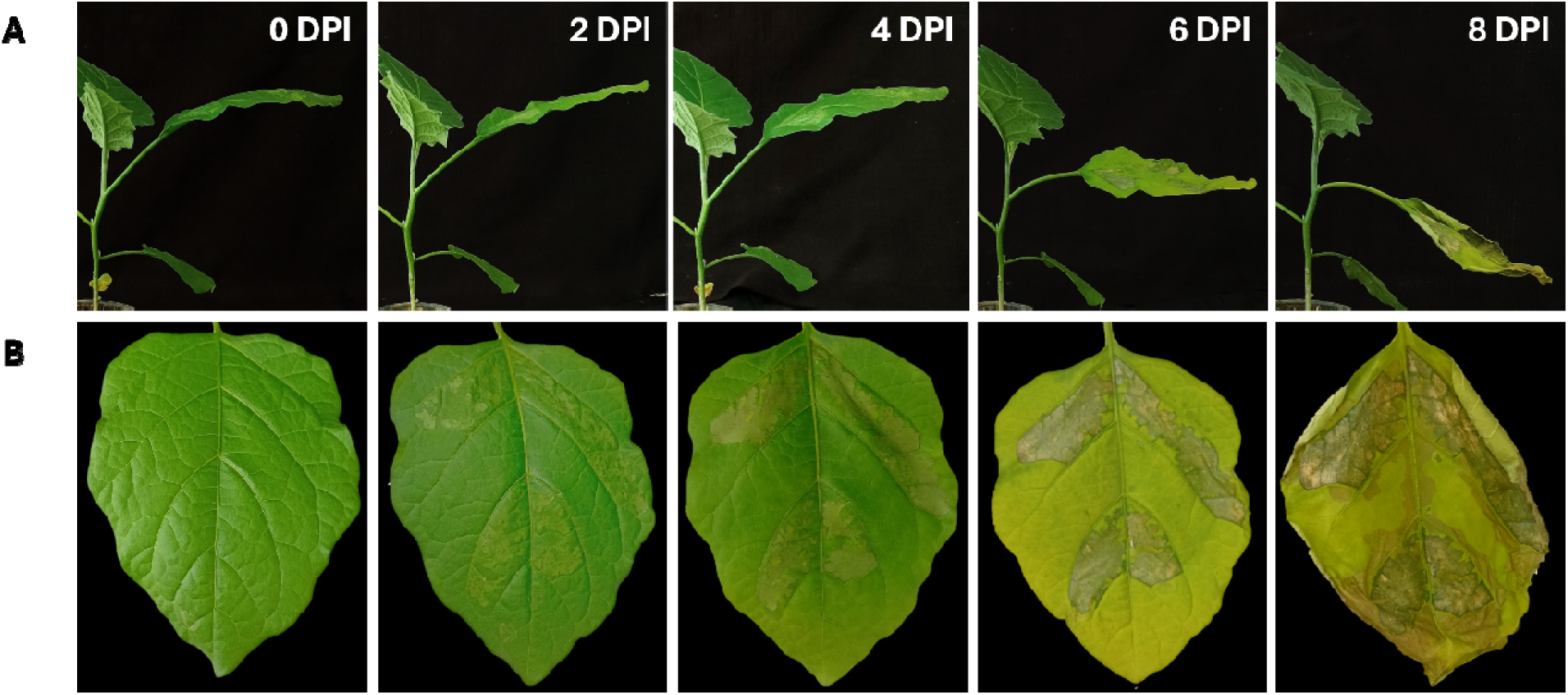
Disease progression in eggplant post-infiltration of *Rps* F1C1 into its leaves. **[A]** Petiole bending; **[B]** Necrotic scar. This experiment was independently performed by four researchers with multiple biological replicates.

### 3.4 Observation of necrotic scars in tomato leaf infiltrated with *Pseudomonas aeruginosa* SPT08, a tomato endophyte

*Pseudomonas aeruginosa* SPT08 was isolated as an endophyte of tomato seedling. This bacterium has been demonstrated to promote plant growth, increase root growth, and protect tomato plants against bacterial wilt when mix inoculated with *Rps* (Giri et al., 2025). It does not elicit hypersensitive response (HR) in tobacco leaves though its genome has type III secretion system (Giri et al., 2025). We infiltrated the SPT08 into tomato leaves. Interestingly, a necrotic scar in the tomato leaflet similar to that of the wild-type F1C1 was observed within 48 h after infiltration with *Pseudomonas aeruginosa* SPT08 (Fig. 4). It is pertinent to note that there was no petiole bending as well as senescence was observed in this case. The scar developed in the cas of *P. aeruginosa* further indicated that leaf infiltration might be a sensitive method for assessing virulence functions.

## 4. Discussion

Inspired by the observation of a hypersensitive response (HR) in tobacco leaves following infiltration with *Rps* F1C1 in our laboratory (Giri et al., 2025), we investigated the effects of leaf infiltration of the pathogen in tomato in the present study. Considering *Rps* is a vascular pathogen, infiltration into the host leaves was performed as an exploratory attempt to examine the plant response. Surprisingly, infiltration induced several distinct patho-phenotypes in tomato plants, including the development of localized necrotic scars that remained confined to the infiltrated area but gradually led to senescence of the infiltrated leaflet (Fig. 3). In addition, infiltration triggered petiole bending and epinasty of the terminal leaflet of the compound leaf (Fig. 5).

Upon infiltration into tomato leaves, this vascular pathogen exhibited distinct virulence-associated phenotypes outside its typical vascular niche. In comparison with the *hrpB::*Ω mutant, which doesn’t result in any scar development (Fig. 4), the results suggest that scar development is likely due to an active interaction between the pathogen and the host, which can be possibly traced directly to the T3S of the pathogen. This interpretation is further supported by the scar formation observed with the *phcA::*Ω, in which the T3S is functional and elicits an HR in tobacco leaves. *phcA* is a master transcription regulator in the pathogen and an established virulent factor (Perrier et al. 2018). The virulence-deficiency of *phcA::*Ω mutant in tomato seedlings is known (Phukan et al., 2019). So, the necrotic scar developed by *phcA::*Ω mutant infiltrated in tomato leaf is indeed surprising. The patho-phenotype developed by *phcA::*Ω mutant in this case indicates that the wilt disease development by the pathogen through systemic infection is more complex than the necrotic lesion formation in the leaf. It is also pertinent to note that the *phcA::*Ω mutant has been proven to be virulent in eggplant seedlings, which is a more susceptible host, by leaf inoculation (Phukan et al. 2019; Kabyashree 2020; Bhuyan et al. 2026). Considering the scar development with *phcA::*Ω mutant, it is proposed that leaf infiltration might be used as a sensitive patho-system to study virulence functions of this pathogen. Necrotic scars were also observed in eggplant leaves following infiltration with *Rps* F1C1 and the *phcA::*Ω mutant, although symptom development was relatively delayed. In contrast, infiltration with *hrpB::*Ω did not induce any visible symptoms in eggplant.

Further, as the necrotic scar remained confined to the infiltrated area, the pathogen likely did not spread beyond the site of infiltration. The characteristic bending at the petiole may represent a plant stress or defense response to the virulence activity of the pathogen within the infiltrated leaflet. A similar response might also be causing epinasty in the terminal leaflet. Together, these observations highlight the ability of the pathogen’s virulence determinants and the plant’s defense response to an immobilized vascular pathogen.

We further investigated leaf infiltration in tomato using *P. aeruginosa* SPT08, a plant growth-promoting endophytic strain previously demonstrated to colonize tomato plants without causing disease and to confer protection from bacterial wilt disease (Giri et al. 2025). SPT08 possesses a functional T3S and exhibits pectinase activity, although no visible cellulase activity has been reported (Giri et al 2025). Interestingly, infiltration with SPT08 resulted in necrotic scar formation in tomato leaves, further supporting the sensitivity of the leaf infiltration assay for studying virulence-associated functions. These observations also reinforce the findings obtained with the *phcA::*Ω mutant, suggesting that vascular wilt disease development is more complex than localized necrotic scarring in the leaf. However, unlike F1C1 and *phcA::*Ω, SPT08 did not induce petiole bending or epinasty of the terminal leaflet. Furthermore, infiltration of SPT08 in eggplant leaves did not result in necrotic scar formation, in contrast to its response in tomato. Future studies involving T3S mutants or other virulence determinants of this bacterium may further establish the utility of this infiltration-based pathosystem for studying bacterial virulence determinants.

It is pertinent to note that the phytopathogen *Xanthomonas oryzae* pv. *oryzae* elicits an HR upon infiltration in tomato leaves, characterized by the appearance of a necrotic scar as a part of the pathogen-non-host interaction (Ray et al., 2000). The necrotic scar in HR also remains limited to the infiltrated area; however, they appear relatively rapidly (within 24-48 h post-infiltration), unlike the necrotic scars observed in the case of the tomato plant (within 48-72 h post-*Rps* infiltration). Also, *Xanthomonas campestris* pv. *vesicatoria* has been reported to elicit HR in tomato leaves within 16 h post-infiltration (Jones and Scott, 1986). Notably, phenotypes such as petiole bending, leaf-epinasty and senescence have not been reported in these HR-associated phenotypes. In a preliminary study from our laboratory (Bhuyan, 2025), *Rps* also elicited HR within 24 h post-infiltration in tobacco (*Nicotiana rustica*) leaves, which, like the eggplant, possess simple leaves; however, the phenotypes of petiole bending and senescence were not observed in the infiltrated leaves in the case of tobacco. These observations suggest that the host plant responses induced by *Rps* infiltration in tomato may be different from the classic HR typically observed in resistant or non-host plants. Another study performed in our laboratory further strengthens that the response induced in tomato plant upon *Rps* infiltration is atypical to the classic HR (Data not presented). It would be interesting to perform a comparative analysis of the two responses in future.

In an earlier study, infiltration in the leaves of tomato, eggplant and bean has been used to study the role of type III effectors in the virulence function of *R. solanacearum* (Macho et al. 2010). However, the study was primarily aimed at comparing the growth of different strains of the pathogen in the leaves using a relatively low pathogen concentration of 10^4^ CFU/mL, and consequently, no visible phenotypic changes could be reported in the infiltrated regions. In contrast, the distinct plant phenotypes observed in the present study highlight the potential of leaf infiltration as a sensitive and underexplored method for investigating virulence functions in Rps.

It is pertinent to note that the development of the leaf-infiltration method by Klement et al. (1964) has proven useful in examining the compatible and incompatible interactions of this broad-host range pathogen with its host plants (Lozano and Sequeira, 1970). In leaves of the tobacco model host (*Nicotiana tabacum* L.), infiltration of incompatible strains of *P. solanacearum* has been reported to induce a Hypersensitive Response (HR). At the same time, infiltration of most compatible strains has been reported to induce a necrotic response that develops relatively slower, allowing the spread of the bacteria to adjoining tissues in the leaf (Granada and Sequeira, 1975). Referring to a compatible or incompatible strain with regard to a given host implies the pathogenicity of that strain in that host, with the former being highly pathogenic and the latter being non-pathogenic or weakly pathogenic in the host (Granada and Sequeira, 1975). More recently, induction of resistance and expression of defense-related genes in tobacco leaves infiltrated with *Ralstonia solanacearum* have been reported using leaf infiltration studies (Kiba et al. 2003).

In conclusion, we propose the phenotypes in tomato leaves observed in this study after infiltration with *Rps* as an easy, consistent, economical, and time-effective approach to assess its virulence functions (Genin 2010; Bhuyan et al. 2024; Bhuyan et al. 2025a; O’Banion et al. 2026). This method effectively reduces any experimental bottleneck that has previously delayed research in bacterial wilt. Beyond these advantages, the development of such virulence assays is essential for facilitating the discovery of newer virulence functions in the pathogen genome (Bhuyan et al. 2026) in the near future.

## Ethics

Our work does not contain experiments involving animals and/or human participants.

## Data accessibility

The article has no additional data.

## Declaration of AI use

We have not used AI-assisted technologies in creating this article.

## Authors’ contributions

M.J. and S.K.: Investigation, methodology, formal analysis, software, validation, visualisation, writing-original draft, writing-review and editing; P.R.D. and Z.R.: Investigation, methodology, visualisation, validation; L.D., S.B. and S.J.G.: Investigation, methodology, visualisation; S.Bhuyan: Investigation, visualisation, software, writing-original draft, writing-review and editing; S.Kushwah and A.K.: Supervision, writing-review and editing; S.K.R.: Conceptualization, supervision, visualisation, writing-original draft, writing-review and editing.

## Acknowledgements

The authors acknowledge Prof. Teresa Ann Coutinho, University of Pretoria, South Africa, for her kind comments that has contributed greatly to the improvement of the manuscript. M.J. and P.R.D. are thankful for the JRF fellowship from CSIR, GoI, New Delhi. S.K. acknowledges UGC, GoI, New Delhi, for the JRF fellowship. Z.R. is thankful to Tezpur University for the institutional fellowship. L.D. is thankful to DST, GoI, New Delhi, for the INSPIRE-JRF/SRF fellowship. S.B. and S.Bhuyan are thankful for the PA-I and RA-I fellowship, respectively, from the DBT, GoI, New Delhi, under grant BT/PR54976/BSA/33/428/2024 awarded to A.K. and S.K.R. S.J.G. is thankful for the JRF fellowship from the DBT, GoI, New Delhi, under grant BT/PR41637/NER/95/1753/2021 awarded to S.K.R. S.Kushwah, A.K., and S.K.R. are thankful to the ANRF-PAIR grant ANRF/PAIR/2025/000029/PAIR.

